# Using changes in host demographic rates to reveal the effects of infection with hidden variable models

**DOI:** 10.1101/2020.02.22.961128

**Authors:** Jake M. Ferguson, Andrea González-González, Johnathan A. Kaiser, Sara M. Winzer, Justin M. Anast, Ben Ridenhour, Tanya A. Miura, Christine E. Parent

**Author notes:** Contributed equally to this paper (Jake M. Ferguson, Andrea González-González).

## Abstract

The impacts of disease on host vital rates can be clearly demonstrated using longitudinal studies, but these studies can be expensive and logistically challenging. We examined the utility of hidden variable models to infer the individual effects of disease, caused by infection, from population-level measurements of survival and fecundity when longitudinal studies are not possible. Our approach seeks to explain temporal changes in population-level vital rates by coupling observed changes in the infection status of individuals to an epidemiological model. We tested the approach using both single and coinfection viral challenge experiments on populations of fruit flies (*Drosophila melanogaster).* Specifically, we determined whether our approach yielded reliable estimates of disease prevalence and of the effects of disease on survival and fecundity rates for treatments of single infections and coinfection. We found two conditions are necessary for reliable estimation. First, diseases must drive detectable changes in vital rates, and second, there must be substantial variation in the degree of prevalence over time. This approach could prove useful for detecting epidemics from public health data in regions where standard surveillance techniques are not available, and in the study of epidemics in wildlife populations, where longitudinal studies can be especially difficult to implement.

## 1. Introduction

The benchmark for detecting the effects of disease on survival and reproduction are longitudinal studies that track individuals through time and can directly link an individual’s infection status to their vital rates. Despite their power, longitudinal studies tend to be relatively rare, especially those concerning wildlife, due to the prohibitive costs and logistics of tracking individuals. Alternatives to longitudinal studies, such as tracking groups instead of individuals, can be used to illustrate differences among populations, however, these comparisons cannot explicitly determine the individual-level impacts of disease. The inability for inferences at one level of aggregation to be valid at a lower level is commonly known as the ecological fallacy.

Hidden variable models, also called latent-variable models, explicitly link processes that are not directly observed to the observed random variables, providing an effective pathway to model complex biological processes with aggregate data (1). Commonly used hidden variable modeling approaches in biology include latent variable regression (2), hidden Markov (3), state-space models (4), and structural equation models (5). In epidemiology, state-space approaches have been successfully used to link the observed number deaths due to disease to the underlying disease dynamics (6) and to distinguish seasonal and epidemic dynamics in public-health surveillance data (7,8). Coupling hidden variable models to inexpensive and commonly collected forms of data, such as population-level measurements of vital rates, could offer a powerful new approach for retroactively determining the individual-level effects of disease on populations. Despite their potential utility, these models have not yet been applied to study the effects of infection on individual fitness components (i.e., survival and reproduction) when individual outcomes cannot be directly observed.

Monitoring vital rates, such as survival and fecundity, is a common approach for managing wildlife populations. Vital rates measurements have been used to provide insights into the past causes of change in population growth (9) and to forecast the effects of potential future environmental conditions on population growth (10). In the context of disease studies, coupling vital rates to disease prevalence has been used to determine how bovine tuberculosis, a mild infection at the individual-level, can lead to large reductions in the population growth of African buffalo (11). More recent work has extended this approach to use vital rate and prevalence measurements to identify the impacts of interactions between multiple diseases (12). Thus, while monitoring vital rates is a potential resource for detecting the effects of disease, the current usefulness of these data is contingent on having detailed individual-level disease prevalence information.

Here, we determined whether the effect of infection on a host’s vital rates could be inferred without direct measurements of infection prevalence. Our approach coupled the observed survival and fecundity rates in a population to a model that predicted the infected status of individuals, the demographic rates of both infected and uninfected individuals and, the probability of obtaining one or more infections. We illustrated these techniques on a tractable experimental host-virus system of *Drosophila melanogaster* and the associated Drosophila C virus (DCV) and Drosophila X virus (DXV) where longitudinal surveys were impossible to perform due to the destructive nature of the sampling. Thus, we were able to measure the change population-level vital rates that occurred due to infection, but we were not able to measure disease incidence directly in our study. Our study reflects a common scenario in wildlife disease ecology where system-specific information on the pathogens is lacking (13). We tested our ability to reconstruct disease dynamics in cases with both single virus infections and coinfections.

## 2. Material and Methods

We used an experimental system to introduce DCV, DXV, and both viruses concurrently in populations of fruit flies. We then used a model to describe the impact of these infections on host vital rates and determined whether we could distinguish the effects of disease on individuals in the population.

### (a) Experimental methods

#### (i) Fly line and husbandry

We used Oregon R wild-type strain of *Drosophila melanogaster* from Carolina Biology Supply. The stock and experimental flies were kept on a 12-hour day/night cycle at 25°C. We housed the stock flies in 25 x 95 mm polystyrene vials (Genesee Scientific) with 10 ml of Jazz-Mix Drosophila Food (Fisher Scientific) prepared following the manufacturer’s instructions. We bleached freshly laid eggs as described in Merkling & van Rij (14) to clear any potential chronic infections before performing the viral challenges.

#### (ii) Infecting pathogens

We used Drosophila C (DCV) and Drosophila X (DXV) viruses as infecting pathogens. DCV has been found in natural populations of at least eight different *Drosophila* species (15,16) and has been extensively used to study and decipher the innate immunity and antiviral responses in the *Drosophila* system (17–21). While it is possible for oral inoculation by DCV to result in systemic infection, it is more often associated with a local immune response and results in a low morbidity and low mortality rates compared to intra-thoracic injections (18,21). DXV is considered to be a cell culture contaminant (22). Although the effects and pathology of DXV are not as well understood as DCV in *Drosophila*, it has been shown that infections by microinjection induce anoxia sensitivity, exhibited by increased rates of mortality when exposed to CO_2_ (22–24).

We used the Charolles strain of DCV isolated from a laboratory population in 1972 (25), and obtained from Dr. Marta L Wayne (University of Florida). We obtained the DXV stock from Dr. Louisa Wu (University of Maryland). Both DCV and DXV were cultured in Schneider’s Drosophila Line 2 (S2 cells) and titrated to a tissue culture 50% infectious dose of 2×10^9^ TCID_50_/ml as previously described (26). Viral stocks were kept in 250 µl aliquots at −80°C.

#### (iii) Oral viral inoculation

We infected flies through an oral inoculation protocol meant to resemble natural routes of infection (14, 27–29). To do so, we collected newly emerged adult flies and left them to age for two or three days, depending on the experiment (survival or fecundity, respectively; see below), and kept males and females in separate vials. On the first day of the experiment, we starved flies for four hours by placing the flies in empty polystyrene vials (30 flies per vial). We then transferred flies to vials supplemented with a 1.91 cm diameter circular piece of Whatman filter paper (Cat No 1001 150) on which 100 µl of viral or mock food mixture was spread. We prepared the viral mixture for single infections by combining 50 µl sucrose 25%, 225 µl DCV or DXV virus stock, 225 µl Schneider’s Drosophila Medium (Genesee Scientific), and 10 µl of edible red dye. The dye allowed us to determine which flies consumed medium and consequently ingested virus. To prepare the coinfection food mix, we added 225 µl of each virus stocks to 50 µl sucrose 25% and 10 µl of red die. For the negative control (i.e., mock), 450 µl of S2 was used in addition to the 50 µl sucrose 25% and 10 µl of edible red dye. We let the flies feed on the food mixture for six hours in the incubator. Flies were sedated on ice and sorted on cold ice packs covered by a Kimwipe. Flies were separated based on whether we could visually detect internal red coloration on the ventral side of their abdomen, indicating recent ingestion of viral medium.

#### (iv) Fecundity experiments

To determine the effects of infection on fly fecundity, we placed a single pair of flies (one male, one female) and incremented the number of pairs up to ten (10 males, 10 females) that had been exposed to virus (indicated by red abdomen) into embryo collection cages (Genesee Scientific, catalog number 59-100). Each collection cage contained a 60 mm grape agar plate (FlyStuff grape agar premix, Genesee Scientific, catalog number 47-102) covered with 150 µl of a yeast mixture made by combining 0.5 g of baker’s yeast and 7 mL of double deionized water. We allowed female flies to lay eggs on each plate for 24 hours before we replaced it with a fresh agar plate with yeast mixture. We conducted this experiment for six days, counting eggs laid on each plate daily using a Nikon SMZ-745 Stereo Zoom Microscope. We allowed the offspring to develop into adults on the plates in an incubator to determine the number of adults that developed from eggs. Agar plates were taped to ensure that offspring could not escape and then frozen for three days after first emergence, at which point adult offspring were counted. We split the breeding pairs into two experimental groups due to space limitations. Odd-numbered populations (1, 3, 5, 7, and 9 mating pairs) were placed in the first group, and even-numbered populations (2, 4, 6, 8, and 10 mating pairs) were placed in the second. We performed two experimental replicates per breeding pair number. Parental flies were collected and frozen at the end of the experiment (day six post-infection), and females were screened for the presence of viral RNA by RT-PCR using fly crude extract (see electronic supplementary material for more details about the RT-PCR protocol and table S1).

#### (v) Survival experiment

To evaluate if a hidden variable model can predict the effects of single and double infections we placed 10 three-day-old flies into fresh food vials in a 1:1 male to female ratio following inoculation. Each treatment (DCV, DXV, and DCV+DXV) and negative control (Mock) consisted of 10 replicate vials for a total of 100 flies per treatment or negative control. We placed experimental vials in the incubator, and counted newly deceased flies in each vial, each day, over 35 days. We also documented the sex of the remaining flies in each vial. Every seven days, adult flies were transferred into new vials with fresh food medium to avoid the emergence of offspring into the experimental population and maintain discrete generations.

### (b) Modeling

Here we describe how we modeled the effects of disease on birth and death rates using hidden variable models. Exposure to these viruses in the experiment does not lead to 100% infection, thus we model each individual’s unobserved infection status as well as the fecundity and survival of individuals in each infected state. Our model then treats the observed population outcomes as a mixture of infected and uninfected individuals.

#### (i) Fecundity models

We tested two models in the fecundity experiment. Our first modeling approach used the observed viral prevalence obtained by RT-PCR to account for the number of infected individuals in the population, while our second modeling approach had the same model structure but did not use any prevalence data. This allowed us to test whether the number of infected individuals in the population could be estimated when that information was not directly observed.

We modeled the time-dependent fecundity of flies in each experimental treatment. For flies not exposed to any virus (hereafter, mock treatment) the rate of offspring production for an uninfected female on day *t* is denoted as *λ*_*U*_(*t*). In the mock treatment, we assumed that the total number of offspring produced on day *t* in a vial with *N*_*U*_(*t*) uninfected females followed a Poisson distribution. A full model specification is given in the electronic supplementary material. For flies that were exposed to DCV or DXV the number of offspring in an experimental vial on day *t* was produced by a mixture of infected and uninfected flies. We estimated the rate of fecundity for an individual fly with either DCV (reproductive rate denoted as *λ*_*C*_) or DXV (reproductive rate denoted as *λ*_*X*_) by modeling the observed number of infected female flies for DCV or DXV and uninfected female flies in a vial, denoted as *N*_*C/X*_(*t*) and *N*_*U*_(*t*), respectively. Similarly, we modeled the number of coinfecteds, *N*_*COI*_, as a mixture of uninfecteds, DCV infecteds, DXV infecteds, and coinfecteds.

We modeled the individual fecundity rates as, 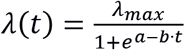, a function that increases over time, denoted as to the asymptotic value *λ*_*max*_. The fecundity at the start of the experiment (*t* = 0) is controlled by the parameter *a*, and the change in fecundity with time is determined by the parameter *b*. Our model assumed that being infected only affects the asymptotic fecundity, *λ*_*max*_, for each experimental treatment and the different values under each treatment are denoted as *λ*_*U*_, *λ*_*C*_, *λ*_*X*_, and *λ*_*COI*_ for the uninfected, DCV, DXV, and DCV+DXV infected flies, respectively.

We also modeled the effects of single infection and coinfection on fly fecundity when the number of flies in each infection class was unobserved. In this scenario, we modeled the number of infected females in a vial as a fixed proportion of the total number of females. For populations exposed to DCV, the number of females infected was *N*_*C*_ =*N* ⋅ *π*_*C*_, where *N* is the known number of individuals in a vial, and *π*_*C*_ is the unknown proportion of flies that are uninfected with DCV. For a population exposed to DXV the number of infected females is *N*_*X*_=*N* ⋅ *π*_*X*_. Finally, for a population exposed to both DCV and DXV we modeled females as infected with DCV but not DXV as *N*_*C,COI*_=*N* ⋅ *π*_*C*_⋅ (1 − *π*_*X*_), females infected with DXV but not DCV as *N*_*X,COI*_=*N* ⋅ *π*_*X*_⋅ (1 − *π*_*C*_), and females coinfected with DCV and DXV as *N*_*COI*_=*N* ⋅ *π*_*C*_⋅ (1 − *π*_*X*_). This model assumed that the probability of becoming coinfected is not affected by previous infections.

We fit all fecundity models in JAGS (30) using an adaptive phase of 10^5^ draws, a burn-in of 10^4^ draws, followed by 10^5^ draws from the posterior. We examined the posteriors graphically for convergence as well as checking the Gelman and Rubin convergence diagnostic (31). For all parameter estimates we report the posterior mode, calculated using the Venter estimator (32), and the standard deviation. We compared parameter estimates of *λ*_*U*_, *λ*_*C*_, *λ*_*X*_, and *λ*_*COI*_ for the models with and without prevalence information to determine whether both modeling approaches yielded consistent answers.

#### (ii) Survival models

We analyzed the effects of DCV, DXV, and coinfection on the mortality rates of flies by treating the experimental data as mixtures of individuals with different possible states (e.g., uninfected or infected). While we knew that the flies ingested the virus in these survival experiments, we were not able to determine which flies cleared the virus and which became diseased. This is because determining the state of infection in flies required destructive sampling, which was problematic for estimating survival. Therefore, our experiments are composed of a mixture of an unknown number of unobserved uninfected and infected individuals.

In the fecundity experiment described above, we assumed, due to the short nature of the study, that infection rates were constant over the 6-day trial. In contrast, we expected the number of uninfected flies in this experiment to change over the course of 35 days. We modeled the population dynamics of the flies and the disease using a susceptible-infected model (SI model) with mortality. We did not include recovery transitions (i.e., transitions of infected individuals to recovered or susceptible) in our model, therefore the only possible transition for infected individuals was death. The transitions used in the single infection and coinfection models are illustrated in figure 1 and the full model specification is provided in the electronic supplementary material.

**Figure 1:**
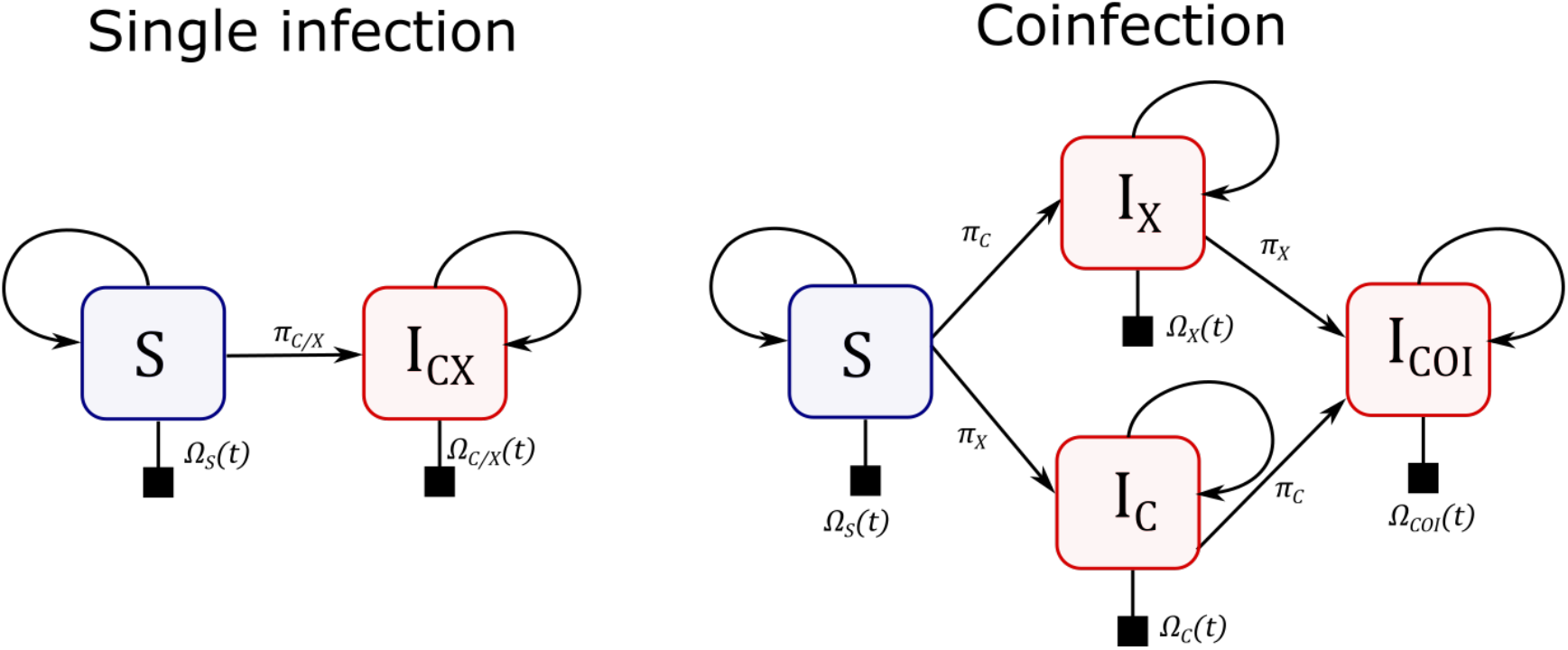
State transitions for an individual with a single disease and multiple diseases (left and right illustration, respectively). Blue boxes are susceptibles (denoted as state S). Red boxes are infecteds with DCV (*I*_*C*_), DXV (*I*_*X*_), or coinfected (*I*_*COI*_). The probability of death of each individual is denoted as *Ω*(*t*), with appropriate subscript, and the probability of a transition from one state to another is denoted as *π*, with an appropriate subscript. For example, the probability of transitioning from susceptible to infected with DCV is *π*_*C*_.

We modeled the probability of survival of an individual to day *t* as 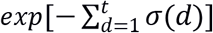 where the discrete hazard function, *σ*(*d*) = *exp*[*α* + *β*_*transfer*_ ⋅ *Transfer* + *β*_*age*_ ⋅ *d*], determines the daily probability of death. The parameter *α* determines the time-independent probability of survival while *β*_*age*_ accounts for the age of the fly (*d* is days since emergence) on survival and *β*_*transfer*_ is the effect of the number of days after the last transfer (*Transfer* is the number of days, from 0 to 6, since the last vial transfer), as described in the experimental methods. These transfers occur each week and we observed increases in fly mortality with time since the last transfer and a corresponding decline once the transfer occurred. This mortality pattern likely occurred due to the medium becoming sticky over time and trapping flies. The expected lifetime of an individual, *L*, can be calculated from the survivorship as 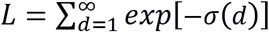.

Our model assumed that all individuals began in the susceptible class (*S*(*t* = 0) = 10) with no individuals in any infected class (*N*_*C*/*X*_(*t* = 0) = 0). Susceptible individuals could become infected with DCV or DXV with the daily probability of infection (*π*_*C*/*X*_), or they could die with the daily probability of death (*Ω*_*S*_(*t*). The daily probabilities of death are denoted as *Ω*(*t*), with a suitable subscript

We assumed that transitions between states were binomially distributed (33). Thus, the number of susceptibles that died between day *t* − 1 and day *t* was binomially distributed with probability *Ω*_*S*_(*t*) and size *S*(*t* − 1). For the single infection model, the total number of infected individuals, *I*_*C/X*_(*t*), on day *t* was the number of susceptibles on the previous day that survived and became infected plus the existing infecteds from the previous day that survived (illustrated in figure 1). For the model with coinfected flies, we included the possibility of single infections by DCV or DXV and the possibility of becoming infected with both viruses. We modeled transitions between states as denoted in figure 1 where the probability of becoming coinfected was assumed to be independent of the current disease state.

We used simulations of the mean-field form of our SI model (34) and Approximate Bayesian Computation (ABC) (35) to obtain parameter estimates. We proposed 10^8^ unique sets of parameters drawn from vague prior distributions (reported in the electronic supplementary material). From these simulations, we selected the top 10^3^ proposals as draws from the posterior. We applied the local-linear regression adjustment of Beaumont et al. (36) to estimate the posterior of the parameter estimates using the Epanechnikov kernel with the smoother-range set to the range of our draws from the posterior. This smoother-range corresponded to a tolerance value for the ABC of *δ* = 6.5.

We were able to calculate the marginal distribution of survival when no disease was present, thus we were able to estimate the parameters *α*_*S*_, *β*_*transfer*_, and *β*_*age*_ using a fully specified model to determine if the ABC model estimates are consistent with the full likelihood. We used JAGS with an adaptive phase of 10^4^ iterations, burn-in of 10^4^ iterations, and 10^4^ draws from the posterior. We used the same priors as the ABC model described above and examined the posteriors graphically for convergence.

## 3. Results

### (a) Fecundity

We found that the per-capita fecundity increased over the first two days of the experiment, reaching an asymptote by the third day of the experiment (figure 2). The estimates of the predicted average fecundity through the course of the experiments are illustrated in the electronic supplementary material figure S1. From here on, we distinguish between the true parameter and estimates of parameters using a circumflex. When using viral prevalence information, we estimated the maximum daily fecundity of an uninfected female (reported as mode (standard deviation)) to be 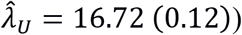. Infected females both had lower fecundity levels with the maximum fecundity of a DXV infected female estimated as 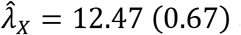 and a DCV infected female as 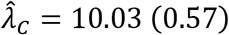. The maximum daily fecundity of coinfected females was less than both DCV and DXV individuals 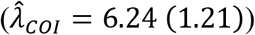. In each comparison the HDI did not overlap 0. We also explored the effects of the total number of adult males and females in each embryo collection cage. While we also found statistically significant effects of both adult male and adult female density on estimates of *λ*, these estimates had little impact on the overall estimates of per-capita fecundity and their interpretation in the context of this study (electronic supplementary material, table S2).

**Figure 2:**
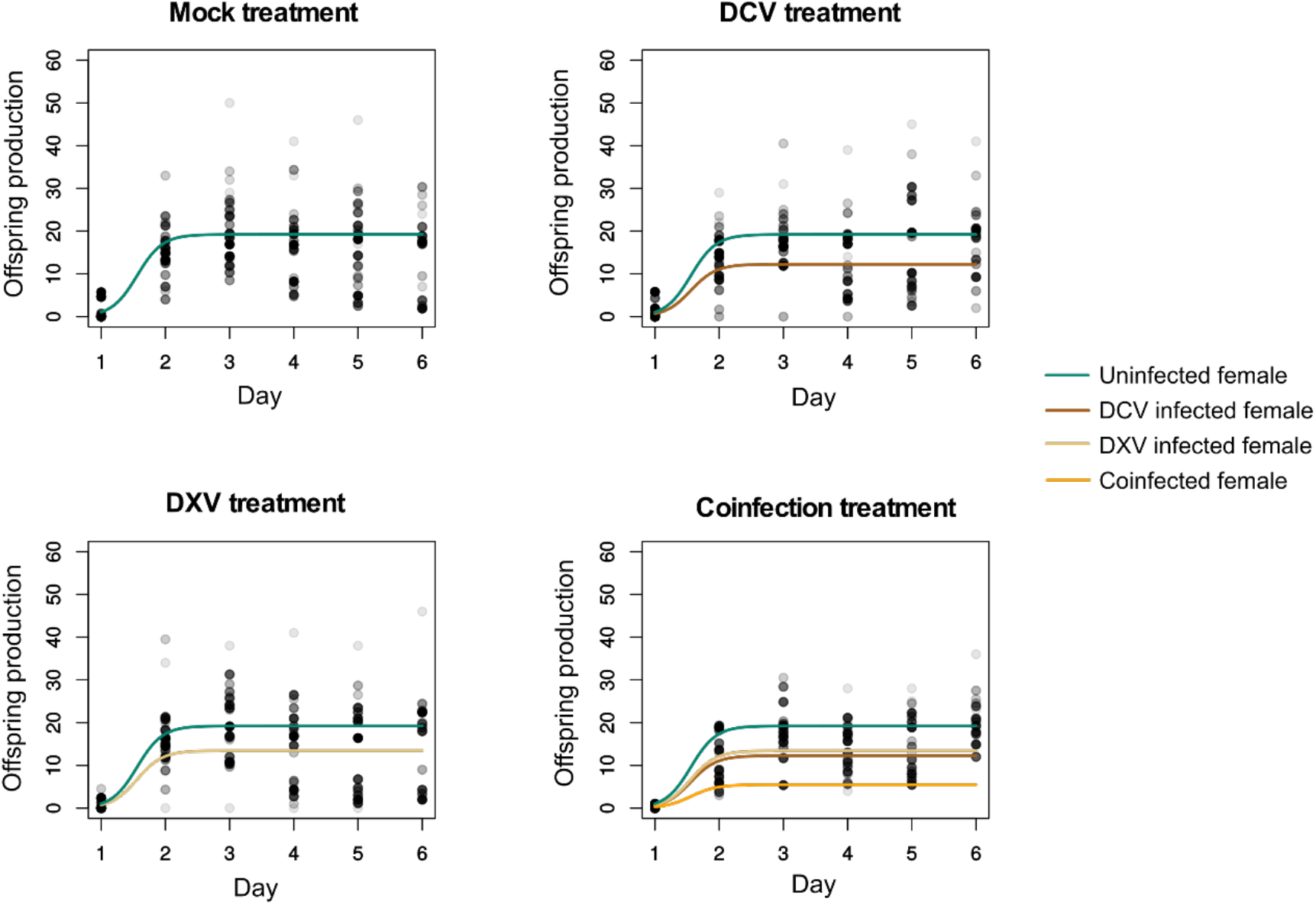
The points are the observed per-capita fecundity data in each experimental vial, shaded to indicate the degree of overlap. Colored lines are the predicted fecundities for an uninfected, infected, or coinfected individual estimated using prevalence information.

When not using viral prevalence information, fecundity estimates were highly uncertain (figure 3). The maximum daily fecundity of a DCV-infected individual was estimated to be 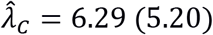. For DXV infected individuals we estimated daily fecundity as 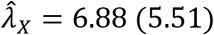 and for a coinfected individual we estimated a maximum fecundity of 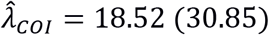. In all cases, estimates of the maximum fecundity of individuals when not using prevalence data were consistent with estimates that did use prevalence data (HDI’s all overlapped 0). Despite the statistical consistency with the estimates using prevalence data, estimates made without prevalence data had such high uncertainty that they were of little practical value (estimates and uncertainties reported in electronic supplementary material, table S3).

**Figure 3:**
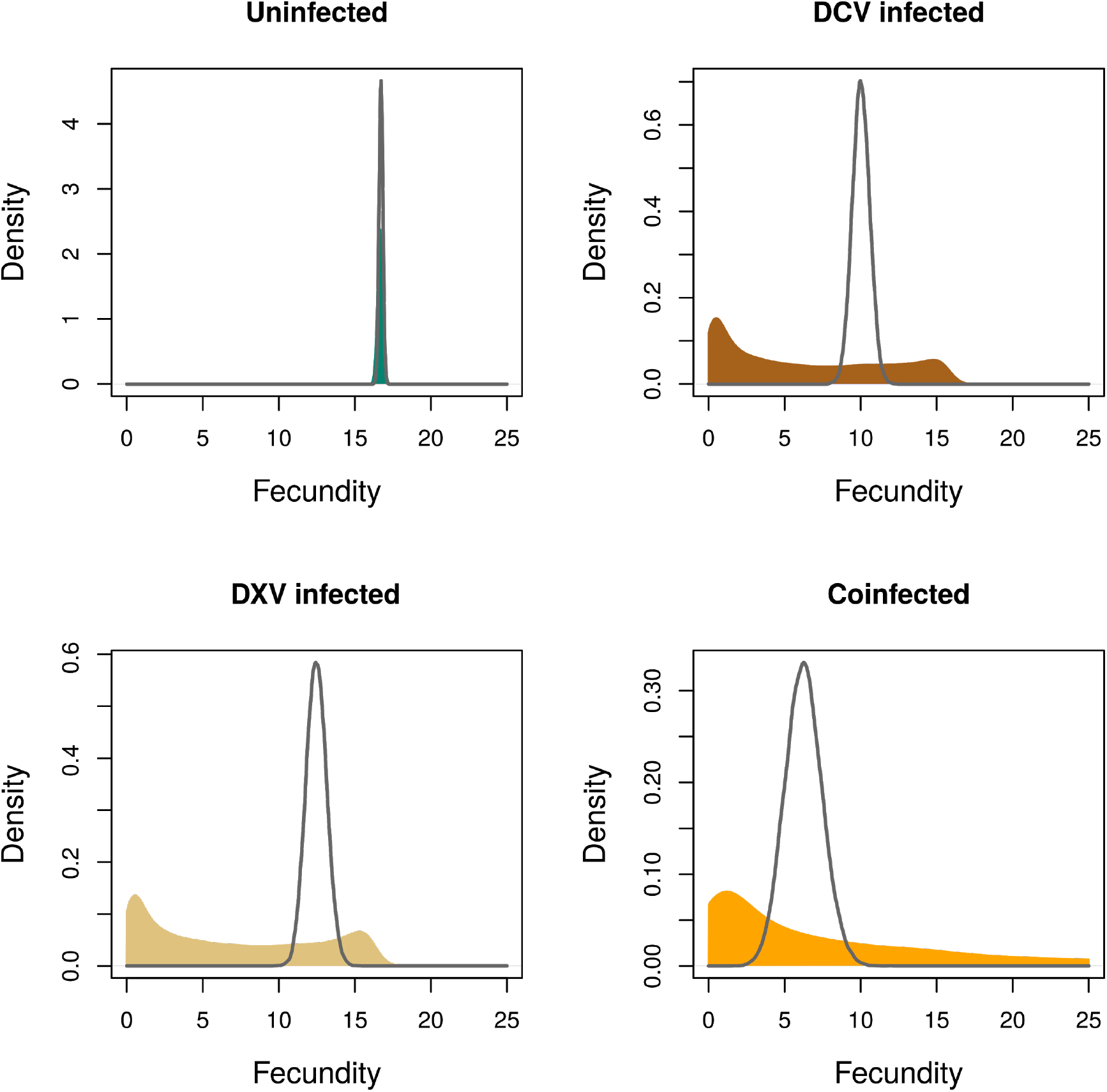
Posterior distributions of the maximum fecundity of a female when disease prevalence is observed (black lines) and when disease prevalence is unobserved (colored area).

We compared the estimated probability of being diseased to the observed prevalence in each vial (electronic supplementary material, figure S2), finding that our estimate of DCV infection, 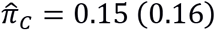, was consistent with the average observed infection probability of 0.07 in the DCV experiment. Our estimate of DXV infection, 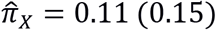, was consistent with the observed infection probability of 0.13 in the DXV experiment, and the estimated coinfection probability, 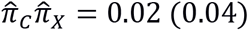, was consistent with the observed coinfection probability of 0.03.

### (b) Survival

By the end of the survival experiment (day 35), mortality rates were 48% in the mock treatment, 60% in the DCV treatment, 67% in the DXV treatment, and 77% in the coinfection treatment. Thus, while survival rates among treatments were similar at the start of the experiment, they differed by the end of the experiment. Treatment-level mortality rates over the course of the experiment are presented in figure 4.

**Figure 4:**
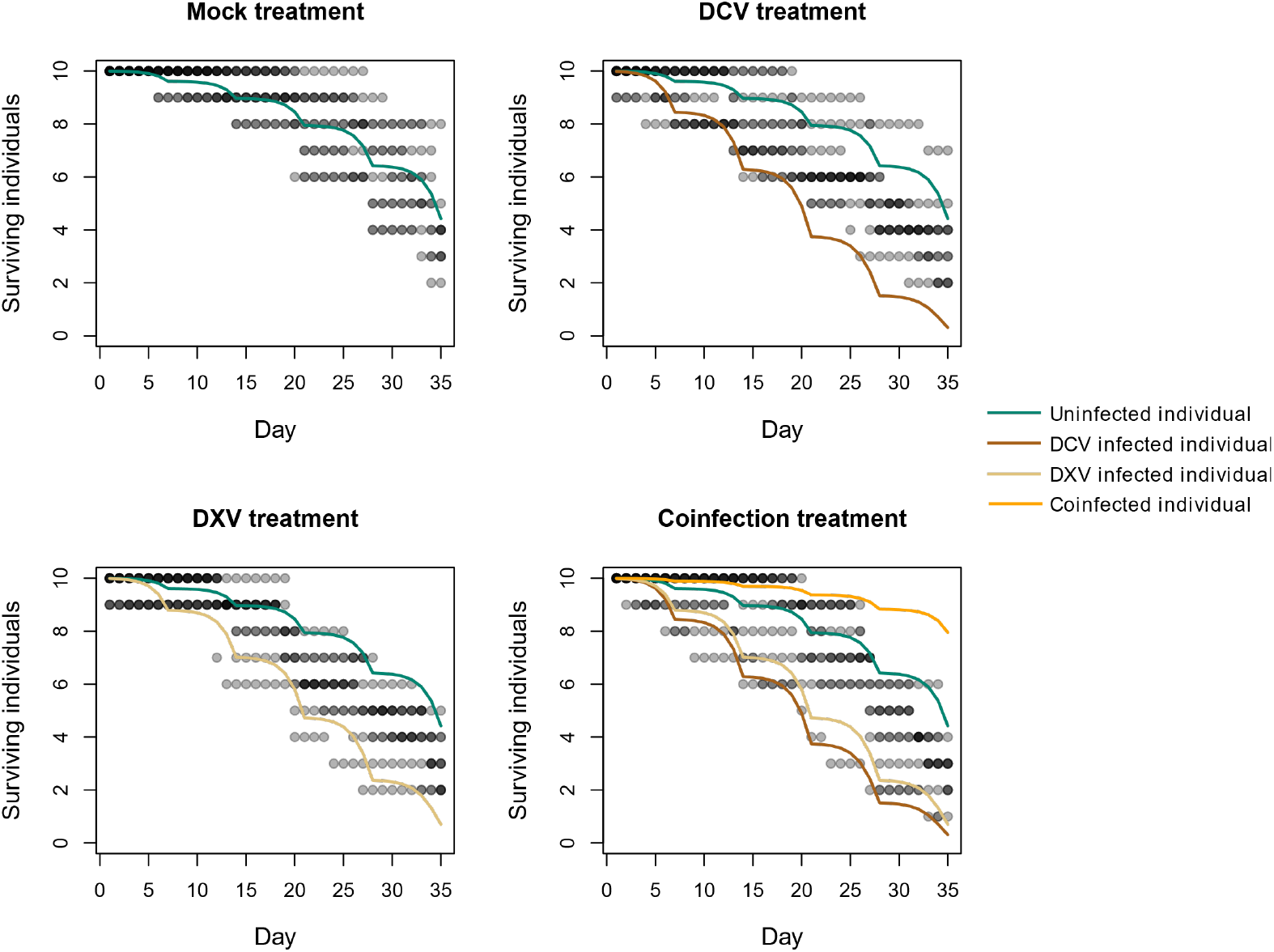
Points are the number of surviving individuals in each vial. Transparency is used to indicate overlapping values. Colored lines are the model predictions for an individual fly that is uninfected, infected, or coinfected.

Our comparison of the ABC estimates and the likelihood estimates indicated that parameter estimates were statistically consistent. From the likelihood model, we estimated the time-independent survival of uninfected flies 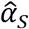 as −8.80 (0.64) (ABC estimate −7.94 (0.94)), the effect of time on fly survival, 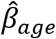, as 0.10 (0.02) (ABC estimate 0.09 (0.01)), and the effect of days since the last vial transfer on survival, 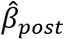, as 0.67 (0.11) (ABC estimate 0.51 (0.17)). In all cases the 95% HDI’s of the difference between posterior approximated by ABC and the full model overlapped zero. Slight differences between these estimates arise because the parameters in the ABC model included additional information through the additional infection treatments that were not able to be accounted for in the likelihood model.

Our posterior estimates indicated that survival decreased with both the time between transfers and age. We found that the survival of a DCV or DXV infected individual was lower than an uninfected individual (95% HDI’s did not overlap 0). The survival of coinfected individuals was highly uncertain, and we found that the HDI’s overlapped 0 for all other pairwise comparisons with the coinfected survival parameter, meaning that we could not statistically distinguish between a coinfected individual and an uninfected, DCV infected, or DXV infected individual (supplementary material, figure S3, table S4). The estimated lifetime of flies that were not subject to the effects of vial transfers (*β*_*transfer*_ = 0) were estimated to fall between 63 and 77 days depending on infection status. The estimated lifetimes of flies infected by DCV or DXV were both lower than uninfected flies (supplementary material, figure S4 left panel, table S4). The mean lifetime of coinfected flies was estimated to be higher than an uninfected individual; however, the high uncertainty in this estimate indicates that it is not a reliable inference.

Our estimates of the probability of infection in the SI model found that the daily probability of becoming infected with DCV 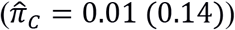 or DXV 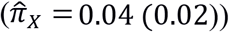 were statistically similar, as the 95% HDI overlapped 0. The probability of being coinfected by DCV and DXV simultaneously on the same day was 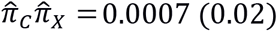 (supplementary material, figure S4 right panel and table S4).

## 4. Discussion

Our results show that hidden variable modeling approaches can be useful for detecting the per-capita effects of disease outbreaks from commonly collected forms of population monitoring data such as survival rates. However, this approach is only suitable for inferring the individual-level effects of infection when two conditions are met. First, the overall effect of infection on the population’s vital rate must be strong enough to detect a statistical difference from an uninfected population. Second, the underlying epidemiological dynamics must generate suitable variation in the degree of prevalence to link changes in the population’s vital rate to the per-capita change in prevalence. These conditions can be met by studying epidemic dynamics over suitable timescales.

Our fecundity experiments did not meet the necessary conditions to identify per-capita disease effects. While we were able to predict these effects very well when prevalence information was available, our analysis failed to accurately estimate prevalence and the effect of disease on fecundity when this prevalence information was removed. Our models were not able to distinguish between very few infected individuals in the population with low fecundity or many individuals with only slight differences from a population of uninfected individuals (figure 3).

Our survival experiment met both conditions necessary to jointly estimate per-capita vital rates and disease prevalence. The increasing discrepancy between experimental treatments and an uninfected population over the course of the experiment allowed the SI model to determine disease prevalence, which in turn allowed us to accurately determine the per-capita effects of disease. While we could detect the effects of single infections on fly survival, we had little ability to detect the effects of coinfection. This limitation arises because the observed outcomes of the coinfection experiment could be adequately explained by either DCV or DXV infections alone. To distinguish the effects of coinfection with these types of data, the population-level effect of coinfection would need to differ from the outcome of singular infections.

We found qualitatively similar effects of DCV infection on both fecundity and mortality (when controlling for the effects of vial transfers on our flies) as past experiments. While we found that fecundity declined with DCV infection, Gupta et al. (37) found that the response to DCV infection depended on host genotype with some genotypes responding to infection with a decrease and others with an increase in fecundity. Previous work on fly survival that has used oral challenge of DCV has found increased mortality soon after exposure (38), and our survival results match the morality rates estimated by Ferreira et al. (18). Although Ferreira and colleagues (18) used a different genetic line (we used Oregon R and they used W^118^) and their infection protocol and dosage differed slightly, they found the same mortality rate of 25% by the end of their 20-day experiment. Additional studies using oral infection have found comparable mortality rates of less than 10% by day 15 or nearly 20% by day 15 post-infection (29). In contrast, Gupta et al. (37) found that inoculation of DCV did not increase mortality relative to unexposed flies, a result that could be explained by their use of a lower viral dosage than the one used in our study or by differences in the density flies were kept at post virus exposure (their flies were kept in isolation).

To our knowledge, our study is the first to determine the effects of oral inoculation with DXV on fly fecundity and survival. We found that both the per-capita effects and the transmission probability of DXV were similar to DCV. While DXV infected flies had a slightly higher estimated fecundity than DCV flies, all other model estimates were statistically indistinguishable between DCV and DXV. We did find that coinfected flies had fecundity levels significantly lower than both DCV and DXV infected flies. These results are consistent with an additive effect of both viruses on fecundity, though the large degree of uncertainty in our estimates also means that a large number of alternative coinfection outcomes are plausible. We could not identify statistical differences between the survival rates of coinfected flies with those infected by DCV or DXV, though we were able to make precise estimates of these parameters. However, our estimate of coinfection on survival rates was highly uncertain.

We showed that hidden variable models can be a useful tool for detecting the effects of per-capita disease from existing monitoring data; however, there are significant limitations when using this indirect approach because we are unable to make direct causal inferences about the effects of disease. In our survival model, we assumed that the probability of acquiring each virus was constant through time and independent of the current number of infected individuals. Retransmission once an individual becomes infected would lead to an additional transmission pathway that depends on the current number of infected individuals in the population. Our fecundity model further assumed that there was no effect of disease on senescence. However, our survival estimates (figure 4) indicated that by the end of the fecundity experiment, we might expect slight differences in fly survival between infected and uninfected individuals.

We made the important simplifying assumption in the coinfection model that infection of diseases occurred independently, that is, the infectious status of an individual did not affect its probability of further infection by the other virus. However, previous work has determined that interactions of disease with the immune system can yield a rich set of epidemiological outcomes. For example, it is well known that coinfection can present interactions that enhance or attenuate the severity of a disease (39,40) or can impact incidence (eg, 41). These effects can also be highly context specific. For example Gonzalez et al. (42) and Marchettoe & Power (43) have shown that the order of infection can govern how diseases impact their host.

While we were able to precisely infer the effects of coinfection in our fecundity experiments when coinfection rates were available, estimation precision was low when these rates were unavailable. The result of our coinfection experiments could be interpreted in several ways because multiple mechanisms could have driven qualitatively similar patterns in our vital rate data (44). Our results were consistent with either the competitive exclusion of DXV by DCV, the competitive exclusion of DVC by DXV, or the coexistence of DCV and DXV. Our inability to distinguish between these outcomes arose due to two factors. First, DCV, DXV, and coinfection had very similar effects on fly survival. Second, the low number of coinfected individuals in our experiment limited our ability to make precise estimates of coinfection effects.

We expect that our hidden variable modeling approach will not only be useful for inferring the per-capita effects of disease from experiments such as ours, but also could be developed into a tool for wildlife health surveillance. While a number of methods exist to detect wildlife epidemics early, many rely on explicitly surveying infection levels (e.g., 45) and may require substantial resources to implement. Our modeling approach could be used to detect emerging epidemics from existing vital rate data that are often collected for management purposes. For example, surveys of European harbor seal mortality detected two anomalous events marked by increased mortality, in 1998 and 2002. These events turned out to be outbreaks of phocine distemper (46). New databases, such as the Marine Mammal Health Monitoring and Analysis Platform (47), have emerged to collect information on mortality events in near real-time. Coupling latent variable models to these data will allow us to potentially explain anomalous mortality events due to disease and forecast the effects of disease outbreak.

There is now a recognition that the complexity of epidemiological processes is much greater than the simple host-disease paradigm. Many epidemiological outcomes are driven by simultaneous infections (48) and across multiple hosts (49). While longitudinal studies are still the gold standard for studying the effects of these diseases on host fitness, the amount effort required to study these emerging frontiers in natural settings necessitates alternative approaches that utilize commonly collected forms of observational data. We expect that hidden variable modeling approaches will help improve the link between modern epidemiological theory and empirical data.

## Supporting information

electronic supplementary material

## Data accessibility

Data and code will be uploaded to Dryad upon acceptance.

## Authors’ contributions

C.E.P. conceived, coordinated and supervised the study. C.E.P., J.A.K, S.M.W. and J.M.A. designed experiments. A.G.G., J.A.K., S.M.W., and J.M.A. performed experiments. T.I.M. provided lab resources, protocols and equipment and co-supervised the study. J.M.F. and B. R. carried out the statistical analyses. J.M.F. and A.G.G. wrote the paper. C.E.P. provided revisions to the manuscript. All authors read and approved the final version of the manuscript for publication.

## Competing interests

The authors declare that they do not have competing interests.

## Funding

J.M.F, A.G.G., J.A.K., S.M.W., J.M.A., B. R., T.I.M. and C.E.P. were supported by the National Institute of General Medical Sciences of the National Institutes of Health under award number P20GM104420. C.E.P. was funded by an Institutional Development Award (IDeA) from the National Institute of General Medical Sciences of the National Institutes of Health under grant number P30GM103324, and by the National Science Foundation under Cooperative Agreement No. DBI-0939454. Additionally, J.M.A. and S.M.W. were supported by the National Science Foundation [grant number DMS-1029485] and undergraduate research awards from the University of Idaho. The content is solely the responsibility of the authors and does not necessarily represent the official views of the National Institutes of Health and of the National Science Foundation.

## Acknowledgements

We thank Ashley DeAguero and LuAnn Scott for their technical assistance. We thank Holly Wichman, and Craig Miller for their helpful feedback on experimental design and data analysis.

