## Supplementary material for "Using changes in host demographic rates to reveal the effects of infection with hidden variable models": electronic supplementary material

### Electronic Supplementary Material for

[*]Contributed equally to this paper

### Experimental methods

#### RT-PCR protocol

We determined whether flies were infected by performing RT-PCR on fly crude extract. We obtained this fly extract by placing individual flies in 2 ml screw-cap tubes with a stainless bead and 50 µl of a 50 mM Tris-Cl, pH 7.5 / 200 µg/ml Proteinase K solution and homogenized them using a TissueLyser for 30 seconds at 30 Hz/sec and incubated the samples for 60 min at 55°C then 5 min at 95°C. We performed RT-PCR on 1 µl of fly extract by using the SuperScript™ III One-Step RT-PCR System with Platinum™ Taq DNA Polymerase (Invitrogen, catalog number 12574-018). Primers listed in Table S1 were used at a concentration of 200 nM each and the following thermal cycling conditions: a reverse transcriptase step for 30 min at 55°C then 2 min at 94°C followed by 40 cycles of 30 sec at 94°C, 30 min at 55°C and 1 min at 68°C, finishing with an extension step of 5 min at 68°C. We ran the RT-PCR products on 1% agarose gels stained with SYBR Safe DNA Gel Stain (Invitrogen) and TAE, and samples that showed either a clear band for DCV (574 bp) or DXV (318 bp) were scored as being infected for the corresponding virus.

Table S1: Set of primers used to determine status infection of individual flies

| Primer name | Sequence (5' - 3') | Reference |
|---|---|---|
| DCV7<br>DCV8 | AGTATGATTTTGATGCAGTTGAATCTC<br>GAAGCACGATACTTCTTCCAAACC | Kapun et al (1) |
| DXV_A2508F<br>DXV_A2825R | TCTCTTATCAACATGGTAGTAAAAAGCG<br>GCCAGTAGGCAATGACTCCTGC | This study |

### Mathematical modeling

#### Prior distributions

##### Fecundity

For the priors of the asymptotic fecundity, $\lambda_U, \lambda_C, \lambda_X$, and $\lambda_{COI}$, we used $\ln \lambda \sim \text{Normal}\left(\mu = 0, \sigma = \sqrt{100}\right)$, $a \sim \text{Normal}\left(0, \sqrt{100}\right)$, and $b \sim \text{Normal}\left(0, \sqrt{100}\right)$. We used uninformative priors for prevalence, $\pi \sim \text{Uniform}(0, 1)$, for the results presented here but also tested moderately informative priors based on past work with DCV (2) with priors of $\text{logit}(\pi_C) \sim \text{Normal}(-1.3, 1)$ and $\text{logit}(\pi_X) \sim \text{Normal}(-1.8, 2)$. On the natural scale these priors correspond to (reported as mode (sd)) a $\pi_C$ of 0.25 (0.17) and for $\pi_X$ of 0.25 (0.26). The lower uncertainty in the prior of $\pi_C$ represents that we know more about DCV than DXV but given the similarities between these viruses we expected to find broadly similar infection rates.

##### Survival

For the time-independent mortality prior distributions we used $\alpha_{S/C/X/CX} \sim \text{Normal}(-8, 5)$, for the probability of infection we used $\pi_{C/X} \sim \text{Beta}(1, 2)$, and for the time-dependent mortality parameters we used $\beta_{transfer} \sim \text{Normal}(0, 1)$ and $\beta_{age} \sim \text{Normal}(0, 1)$.

#### Model specification

##### Fecundity

We assume that for $N_U(t)$ uninfected females at time $t$, each with reproduction rate $\lambda_U(t)$, the number of offspring follows a Poisson distribution, $F_U(t) \sim \text{Poisson}(N_U(t)\, \lambda_U(t))$ The total fecundity in a vial of individuals exposed to DCV or DXV is modeled as a mixture of uninfected and infected individuals as $F_{C/X} \sim \text{Poisson}\left(N_U \lambda_U + N_{C/X} \lambda_{C/X}\right)$, where we have dropped the time argument for readability and used the fact that the distribution of a sum of Poisson random variables with different rates is a new Poisson with a rate given by the sum of individual rates (3). For the coinfection experiment, the mixture of uninfected and infected individuals is

$F_{COI} \sim \text{Poisson}(N_U\lambda_U + N_C\lambda_C + N_X\lambda_X + N_{COI}\lambda_{COI})$, where $N_{COI}$ is the number of coinfected individuals and $\lambda_{COI}$ is the coinfected reproductive rates.

#### Survival

The daily probability of death, $\Omega(t)$, is the probability of dying between day $t$-1 and day $t$, given that the individual survived up to day $t$-1. The time-independent mortality terms for uninfected susceptibles was denoted as $\alpha_S$, and infecteds are hereafter denoted as $\alpha_C$ (for DCV-infecteds), $\alpha_X$ (for DXV-infecteds), or $\alpha_{COI}$ (for coinfecteds). The regression parameters $\beta_{transfer}$ and $\beta_{age}$ were assumed to be independent of infection status.

The total number of infected individuals, $I_{C/X}(t)$, on day $t$ was the number of susceptibles on the previous day that survived and became infected plus the existing infecteds from the previous day that survived. Thus, the expected numbers of susceptible and infected individuals in a population exposed to a single virus given the previous day's population are:

$$E[S(t)] = (1 - \Omega_S(t)) \cdot S(t-1) \; - \; \pi_{C/X}(1 - \Omega_S(t)) \cdot S(t-1)$$
$$E[I_{C/X}(t)] = (1 - \Omega_{C/X}(t)) \cdot I_{C/X}(t-1) \; + \; \pi_{C/X}(1 - \Omega_S(t)) \cdot S(t-1).$$

A description of the expected numbers of individuals in each state for the model with coinfection is:

$$E[S(t)] = (1 - \Omega_S(t)) \cdot S(t-1) \; - \; (\pi_C \; + \; \pi_X \; - \; \pi_C\pi_X)(1 - \Omega_S(t)) \cdot S(t-1)$$
$$E[I_C(t)] = \; \pi_C(1 - \pi_X)(1 - \Omega_S(t)) \cdot S(t-1) \; + \; (1 - \pi_X)(1 - \Omega_C(t))I_C(t)$$
$$E[I_X(t)] = \; \pi_X(1 - \pi_C)(1 - \Omega_S(t)) \cdot S(t-1) \; + \; (1 - \pi_C)(1 - \Omega_X(t))I_X(t)$$
$$E[I_{COI}(t)] = \pi_X(1 - \Omega_C(t))I_C(t) + \pi_C(1 - \Omega_X(t))I_X(t) \; + \; (1 - \Omega_{COI}(t))I_{COI}(t).$$

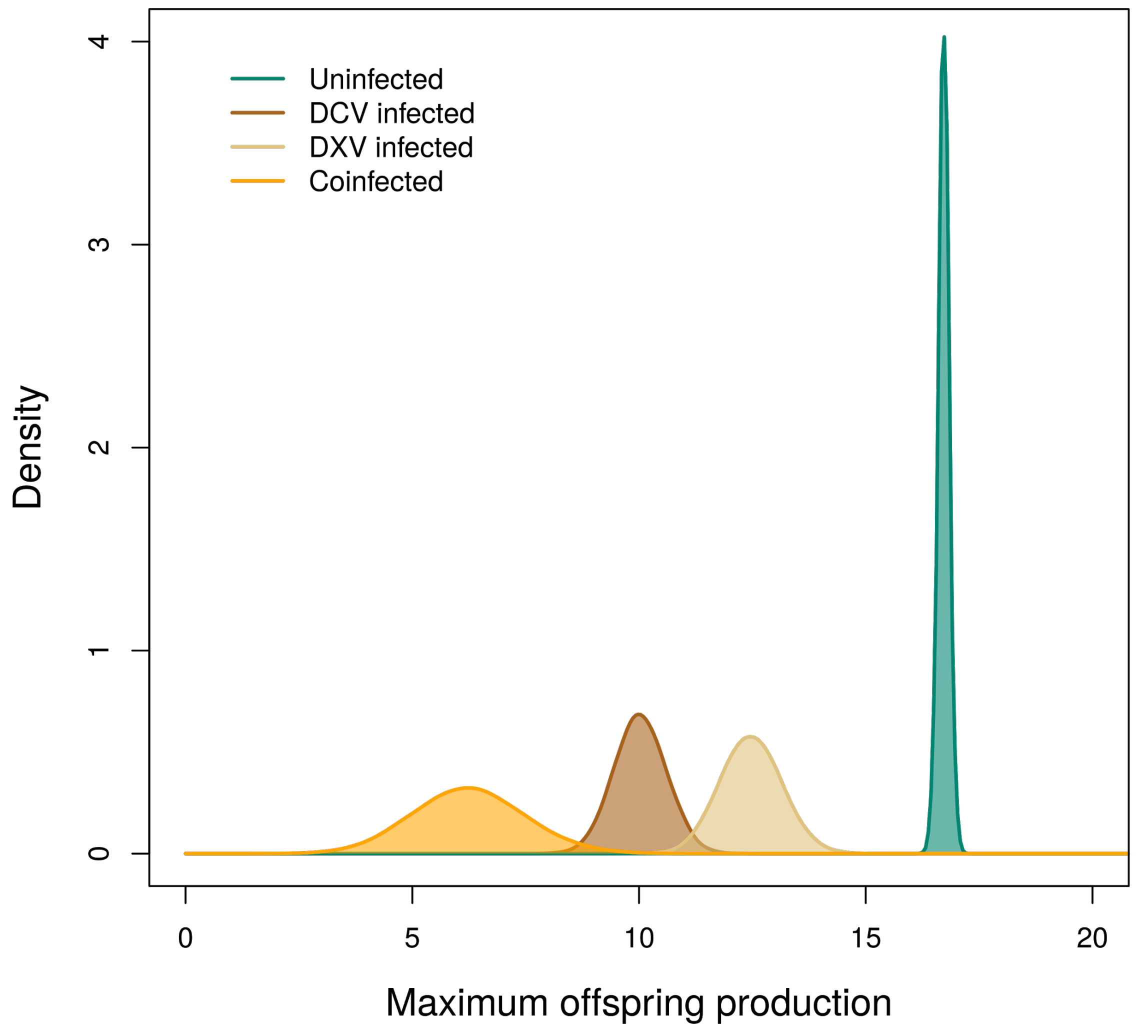


Figure S1: Posterior distributions of the maximum fecundity of a female ($\hat{\lambda}$) when disease prevalence is observed.

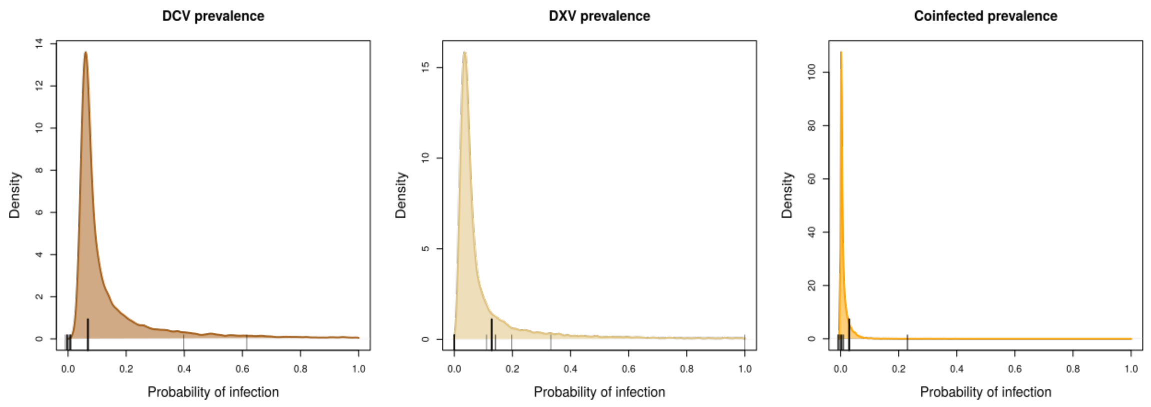


Figure S2: Posterior distributions of the probability of infection for the DCV, DXV, and coinfection fecundity experiments. The rug ticks at the bottom of each plot are the observed prevalence's measured by RT-PCR in each vial on day 6 of the experiment (jittered for clarity). The large tick is the observed mean prevalence.

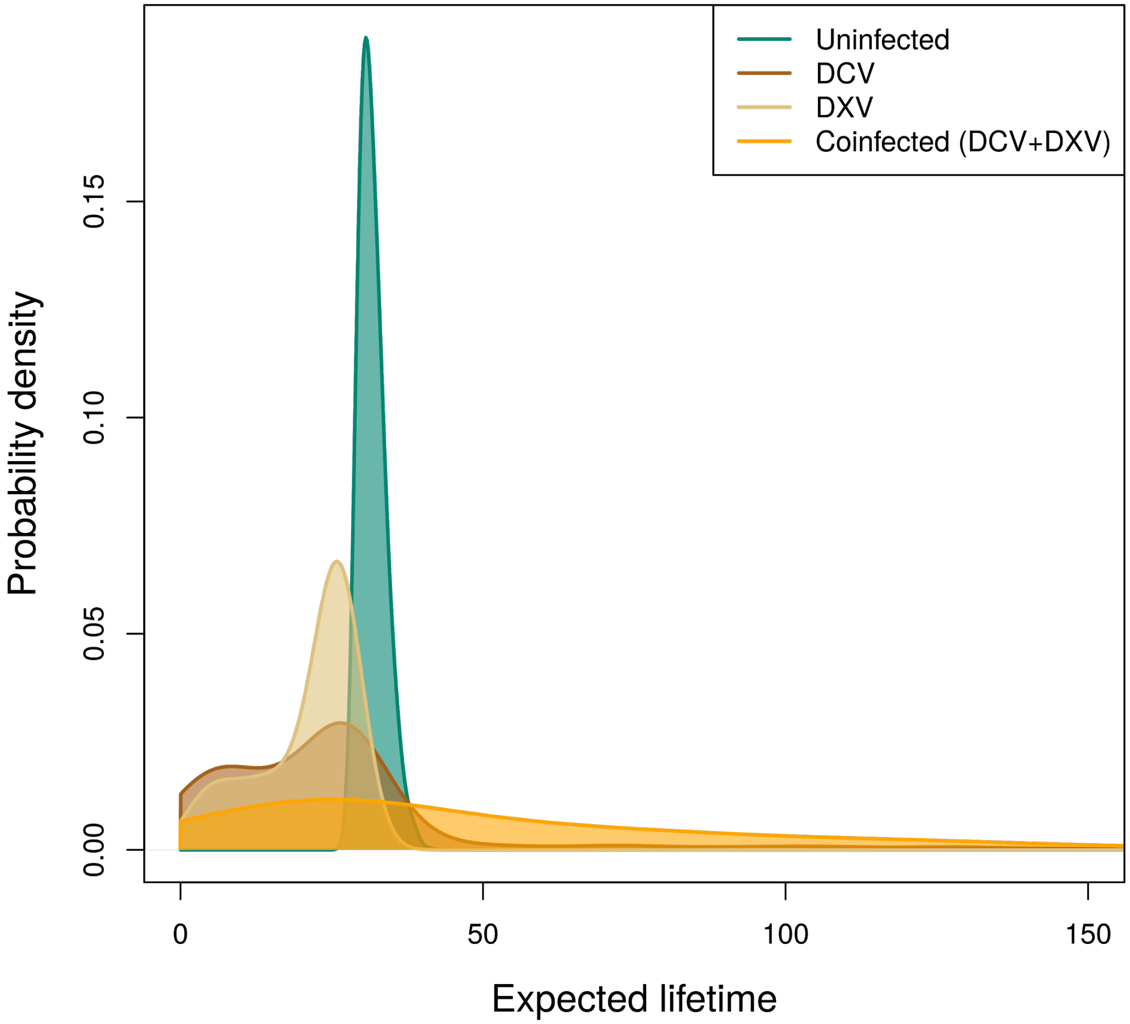


Figure S3: Posterior distributions of the expected lifetime of a fly in a vial with the effects of vial transfers.

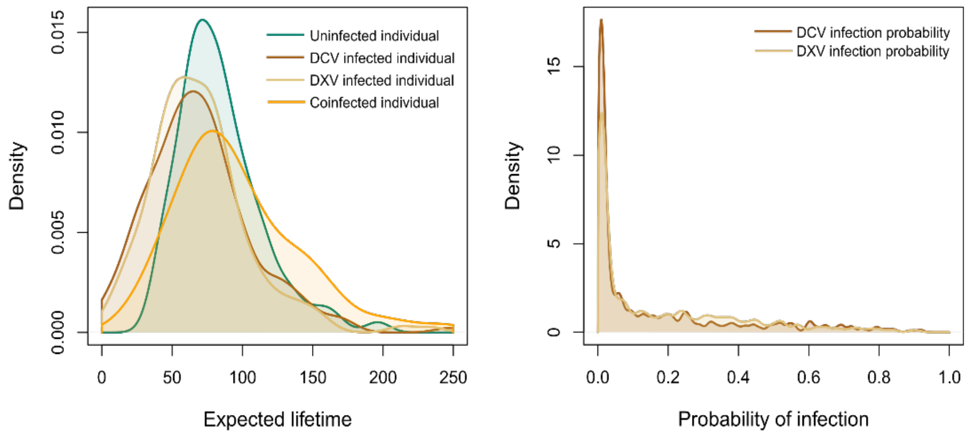


Figure S4: Posterior distributions of the predicted lifetime of a fly in a vial without the effects of vial transfers (left panel), and the daily probability of infection (right panel).

Table S2: Parameters and their estimates in the fecundity model when including the effects of males and female density as covariates.

| Variable | Description | Estimation without prevalence mode (SD) |
|---|---|---|
| $a$ | Determines a fly's initial fecundity | 2.87 (0.06) |
| $b$ | The rate of change in fecundity with time | 4.93 (0.14) |
| $\beta_F$ | Effect of number of females on maximum fecundity | 0.04 (0.01) |
| $\beta_M$ | Effect of number of males on maximum fecundity | -0.06 (0.01) |
| $\hat{\lambda}_U$ | Maximum fecundity of an uninfected individual | 19.94 (0.40) |
| $\hat{\lambda}_C$ | Maximum fecundity of a DCV infected individual | 9.13 (6.44) |
| $\hat{\lambda}_X$ | Maximum fecundity of a DXV infected individual | 7.79 (6.37) |
| $\hat{\lambda}_{COI}$ | Maximum fecundity of a coinfected Individual | 17.79 (30.03) |
| $\hat{\pi}_C$ | Probability of being infected by DCV | 0.15 (0.15) |
| $\hat{\pi}_X$ | Probability of being infected by DXV | 0.13 (0.14) |
| $\hat{\pi}_C\hat{\pi}_X$ | Probability of being infected by both DCV and DXV | 0.03 (0.05) |

Table S3: Parameters and their estimates in the fecundity model estimated with and without viral prevalence information.

| Variable | Description | Estimation with prevalence mode (SD) | Estimation without prevalence mode (SD) |
|---|---|---|---|
| $a$ | Determines a fly's initial fecundity | 2.85 (0.06) | 2.87 (0.06) |
| $b$ | The rate of change in fecundity with time | 5.03 (0.16) | 4.94 (0.14) |
| $\hat{\lambda}_U$ | Maximum fecundity of an uninfected individual | 16.72 (0.12) | 16.67 (0.18) |
| $\hat{\lambda}_C$ | Maximum fecundity of a DCV infected individual | 10.03 (0.57) | 6.29 (5.20) |
| $\hat{\lambda}_X$ | Maximum fecundity of a DXV infected individual | 12.47 (0.67) | 6.88 (5.51) |
| $\hat{\lambda}_{COI}$ | Maximum fecundity of a coinfected individual | 6.24 (1.21) | 18.52 (30.85) |
| $\hat{\pi}_C$ | Probability of being infected by DCV | 0.07 (0.05) | 0.15 (0.16) |
| $\hat{\pi}_X$ | Probability of being infected by DXV | 0.13 (0.07) | 0.11 (0.15) |
| $\hat{\pi}_C\hat{\pi}_X$ | Probability of being infected by both DCV and DXV | 0.03 (0.02) | 0.02 (0.04) |

Table S4: Parameter definitions and estimates for the survival model.

| **Variable** | **Description** | **Estimated mode (SD)** |
| --- | --- | --- |
| $L_S$ | The expected lifetime of a symptomatic fly without the effect of vial transfers | 73.2 (15.7) |
| $L_C$ | The expected lifetime of a DCV infected fly without the effect of vial transfers | 63.3 (19.4) |
| $L_X$ | The expected lifetime of a DXV infected fly without the effect of vial transfers | 64.7 (18.0) |
| $L_{COI}$ | The expected lifetime of a coinfected fly without the effect of vial transfers | 76.9 (51.1) |
| $\beta_{transfer}$ | The effects of days since the last vial transfer on fly survival | 0.51 (0.17) |
| $\beta_{age}$ | The effect of time on fly survival | 0.09 (0.01) |
| $\alpha_S$ | Time-independent probability of survival of symptomatic flies | -7.94 (0.94) |
| $\alpha_C$ | Time-independent probability of survival of DCV symptomatic flies. | -6.50 (1.62) |
| $\alpha_X$ | Time-independent probability of survival of DXV symptomatic flies. | -7.59 (1.41) |
| $\alpha_{COI}$ | Time-independent probability of survival of coinfected flies | -8.44 (3.93) |
| $\hat{\pi}_C$ | Daily probability that an individual becomes infected with DCV. | 0.01 (0.14) |
| $\hat{\pi}_X$ | Daily probability that an individual becomes infected with DXV. | 0.02 (0.18) |
| $\hat{\pi}_C\hat{\pi}_X$ | Daily probability that a susceptible individual becomes infected with both DCV and DXV. | 0.0007 (0.02) |
